# Aerial root formation in Oaxacan maize (*Zea mays*) landraces persists into the adult phase and is minimally affected by soil nitrogen and ambient humidity

**DOI:** 10.1101/2024.10.21.619439

**Authors:** Jennifer Wilker, Rafael E. Venado, Valentina Infante, Caitlin McLimans, Fletcher Robbins, Courtney Phillips, Claudia Irene Calderón, Jason Wallace, Jean-Michel Ané

## Abstract

Maize (*Zea mays*) is the most widely produced crop in the world, and conventional production requires significant amounts of synthetic nitrogen fertilizer, which has negative economic and environmental consequences. Maize landraces from Oaxaca, Mexico, can acquire nitrogen from nitrogen-fixing bacteria that live in a mucilage secreted by aerial nodal roots. The development of these nodal roots is a characteristic traditionally associated with the juvenile vegetative stage of maize plants. However, mature Oaxacan landraces develop many more nodes with aerial roots than commercial maize varieties. Our study shows that Oaxacan landraces develop aerial roots during both the juvenile and adult vegetative phases and even during early flowering under greenhouse and field conditions. Surprisingly, the development of these roots was only minimally affected by soil nitrogen and ambient humidity. These findings are an important first step in developing maize varieties that can reduce fertilizer needs in maize production across different environmental conditions.

## INTRODUCTION

Plant roots support plant growth by fulfilling essential water and nutrient acquisition and anchorage functions. Roots release a significant amount of the plant photosynthates in the surrounding soil, allowing plants to shape their rhizosphere microbiome (Pantigoso et al., 2022; Santangeli et al., 2024). Roots are also the site of intimate associations with symbiotic microbes such as nitrogen-fixing bacteria (diazotrophs) or mycorrhizal fungi that can improve plant nutrition and stress tolerance. Roots predominantly arise from the root apical meristem (RAM), yet in monocots, the shoot apical meristem (SAM) initiates the development of above-ground structures, such as leaves, tassels, and nodal roots (Thompson et al., 2015). Regardless of their origin, these meristems serve as reservoirs of stem cells, perpetually generating new cells to sustain growth and differentiation (Benfey and Scheres, 2000).

Root systems differ vastly among plants, and monocots exhibit a complex root structure distinct from eudicots. The monocot primary root undergoes decay and is rapidly substituted by a fibrous root system composed of adventitious nodal roots, also known as crown roots, originating at the stem base (Hochholdinger et al., 2004; Blizard and Sparks, 2020). Maize (*Zea mays* L.), particularly, features crown and above-ground nodal roots, often referred to as “brace roots,” that reach the ground and serve essential anchorage functions (Hostetler et al., 2021). However, some maize accessions also produce many nodal roots that never reach the ground (hereon referred to as “aerial roots”), and their function remained enigmatic for a long time (**Figure 1A**). In 2018, we reported that landraces from Southern Mexico develop many more aerial roots than other widespread maize varieties (**Figure 1B**) (Van Deynze et al., 2018). We also showed that these maize landraces can acquire large amounts of nitrogen from diazotrophs that live in a gel/mucilage produced by their aerial roots after rain (Van Deynze et al., 2018; Bennett et al., 2020; Pankievicz et al., 2022). Both brace and aerial roots originate from stem nodes derived from the SAM. This developmental process involves four stages: induction, where founder cells acquire the ability to divide; initiation, marked by the observable presence of root primordia in the node; emergence, during which the root primordium elongates; and growth, where elongation continues until the complete disappearance of the root cap (Itoh et al., 2005).

**Figure 1.**
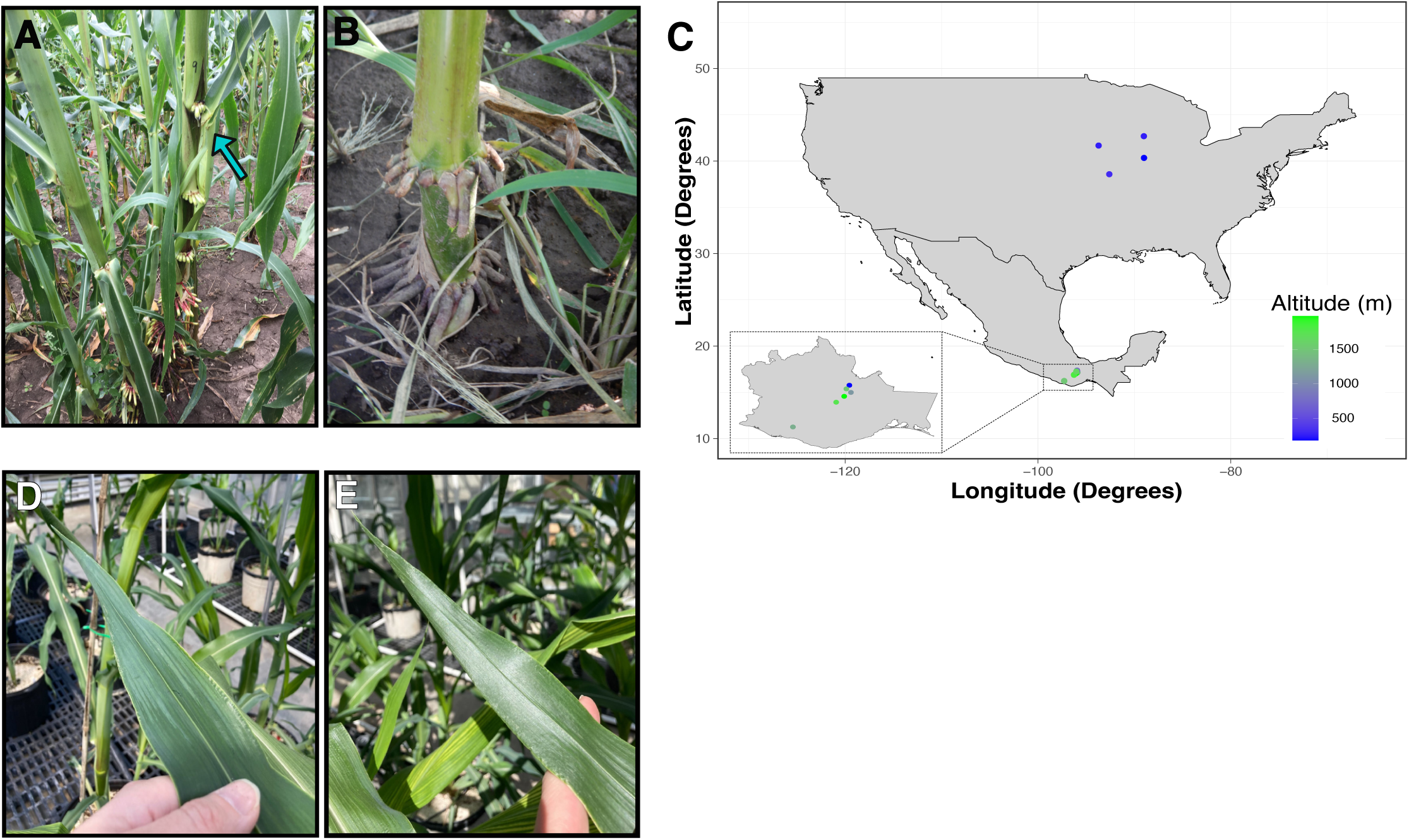
Maize landraces and exPVP lines and their phenotypes. A) Maize landraces and their aerial roots. The blue arrow shows the production of mucilage. B) exPVP PHP02 and its aerial roots. C) Geographical distribution (latitude, longitude, and altitude) of the maize genotypes used in this study. D) Last leaf with epicuticular wax. E) First leaf without epicuticular wax.

The root development process is intricate and dynamic, influenced by genetic and environmental factors. In maize, 21 distinct genes have been identified to affect root development to varying degrees, as reviewed previously (Hostetler et al., 2021). Environmental factors such as water availability, temperature, microbes, and drought often shape root development (Lynch et al., 2012). Some of these factors modulate plant hormones, significantly impacting root development. For instance, waterlogged conditions prompt the development of adventitious nodal roots, facilitated by the accumulation of auxin at the stem base. In contrast, drought stress involves precisely regulating abscisic acid (ABA) and auxin levels to sustain root elongation (Kazan, 2013). Plant hormones play a fundamental role in root development, and their interplay and accumulation affect different root types. Specific hormones, such as auxin and ethylene, positively influence nodal roots, while others, like cytokinin, act as antagonists, hindering their development, as reviewed previously (Singh et al., 2023). Nutrient levels also affect the root system. For example, maize exhibits increased lateral root growth under short-term low nitrogen levels, but prolonged deficiency hampers lateral root development (Gaudin et al., 2011; Sun et al., 2021).

Root development often correlates with other developmental transitions. In maize, brace and aerial root development have been traditionally associated with the juvenile vegetative stage of maize growth (Poethig, 1990; Hostetler et al., 2021). Above-ground nodal roots are sometimes observed at the adult stage but only after lodging (Sparks, 2023). However, Oaxacan landraces develop many more nodes with aerial roots than commercial maize accessions and without lodging (Van Deynze et al., 2018). This observation suggests that these landraces may continue to form aerial roots after the juvenile stage or that the juvenile stage in these accessions is extended as in the *corngrass1* (*cg1*) mutant (Chuck et al., 2007). In maize, the transition from juvenile-adult vegetative phase shift is marked by the disappearance of leaf epicuticular wax, leaf hairs, cell wall composition, and insect and rust resistance changes. Leaves five or six possess epidermal cells responsible for wax production. However, a transition occurs beyond the sixth leaf, marked by differentiation in cell types, notably the development of bulliform cells and leaf hairs. (Freeling and Lane, 1994; Moose and Sisco, 1994; Bongard-Pierce et al., 1996; Sylvester et al., 2001; Poethig, 2003; Riedeman et al., 2008; Riedeman and Tracy, 2010).

Here, we investigated whether Oaxacan landraces produce aerial roots during adult growth under greenhouse and field conditions. In field trials in Wisconsin and Georgia, we examined how environmental conditions influence the formation of aerial roots. Concurrently, we investigated the impact of environmental factors on various traits of aerial roots. Oaxacan landraces were cultivated under three nitrogen levels in one experiment, and in another experiment, they were subjected to differing relative humidity levels. Both experiments took place in greenhouse settings. It is crucial to comprehend the diverse factors contributing to the development of aerial roots if we aim to harness the untapped potential of these landraces in breeding initiatives to reduce nitrogen fertilizers in agriculture.

## RESULTS

### Oaxacan maize landraces develop elongated aerial roots during the adult vegetative and reproductive phases in controlled greenhouse conditions

A greenhouse study was first conducted to examine developmental growth phases and aerial root formation in two inbreds (PHZ51, PHP02) with expired Plant Variety Protection Act certificates (exPVP), one giant heirloom (Hickory King), and three Oaxacan landrace accessions (Oaxa233, Oaxa524, Oaxa733) under controlled conditions. Landraces were obtained from the International Maize and Wheat Improvement Center (Spanish acronym CIMMYT), previously collected from southern Mexico, and the exPVP and giant heirloom were acquired from the American Germplasm Resources Information Network (GRIN) (**Figure 1C** and **Supplementary Table 1**). We used leaf epicuticular wax production as a morphological marker to determine the transition between the juvenile to adult vegetative phase (**Figure 1D and E**) (Freeling and Lane, 1994; Bongard-Pierce et al., 1996). We counted the number of leaves that displayed the presence of wax on the abaxial side of the leaf. The wax appears as an opaque layer on the leaf surface extending from the leaf tip towards the base. Growth of the last leaf with epicuticular wax was significantly different (p = 0.004) among genotypes (**Figure 2A**). PHZ51, PHP02, and Oaxa524 transitioned to the adult phase after the growth of leaf nine, whereas Hickory King, Oaxa233, and Oaxa733, on average, transitioned one to two leaves later. The exPVP genotypes generally exhibited narrower variability for this trait. In contrast, the heirloom and landrace genotypes showed wider-ranging values, probably because the exPVP lines are genetically uniform inbreds, while the heirloom and landraces are heterogeneous and outbred. Anthesis (male flowering) marks the transition from the adult vegetative to the adult reproductive phase. Under greenhouse conditions (12-hour photoperiod), the number of days to anthesis was significantly different among genotypes (p = 0.0012), with the exPVPs and Hickory King tasseling earlier than the landrace genotypes (**Figure 2B**). PHZ51 and PHP02 reached anthesis in less than 70 days, Hickory King in an average of 85 days, and the landraces in over 95 days post-planting. Oaxa233 appears to be photoperiod sensitive, and only one replicate produced a mature tassel. At anthesis, the number of nodes with aerial roots was quantified. The exPVP and heirloom genotypes produced significantly fewer nodes with aerial roots than landrace accessions (**Figure 2C**). For example, PHZ51 produced significantly fewer nodes with aerial roots than Oaxa524 (p = 0.018) and Oaxa233 (p = 0.035). In addition, the adult vegetative phase (the time between the formation of the last leaf with epicuticular wax and anthesis) was longer in the landrace accessions than in the exPVP and heirloom genotypes (Supplementary Figure 1). Potential correlations were investigated among the number of nodes with aerial roots, days to anthesis, and the last waxy leaves. Our analysis revealed positive correlations among all traits in the landraces (**Supplementary Figure 2A**). Specifically, we observed a moderate to high correlation between days to anthesis and the number of nodes with aerial roots (r = 0.66). In contrast, the correlations between the other pairs were very low (**Supplementary Figure 2B**). The number of nodes with aerial roots was also assessed at various intervals between 64 and 135 days after planting (DAP) (**Figure 3**). PHP02 and PHZ51 grew no additional nodes after tasseling (average 70 DAP) and averaged three nodes with aerial roots in both exPVPs (**Figure 3** and **Supplementary Figure 1**). In contrast, the larger Hickory King variety flowered at an average of 85 DAP and averaged five nodes with aerial roots per plant. Intriguingly, the landraces continued producing aerial roots even after anthesis, typically after 100 DAP (**Figure 3** and **Supplementary Figure 1**). These findings suggest landraces continue developing aerial roots into the adult reproductive stage, unlike the other maize varieties.

**Figure 2.**
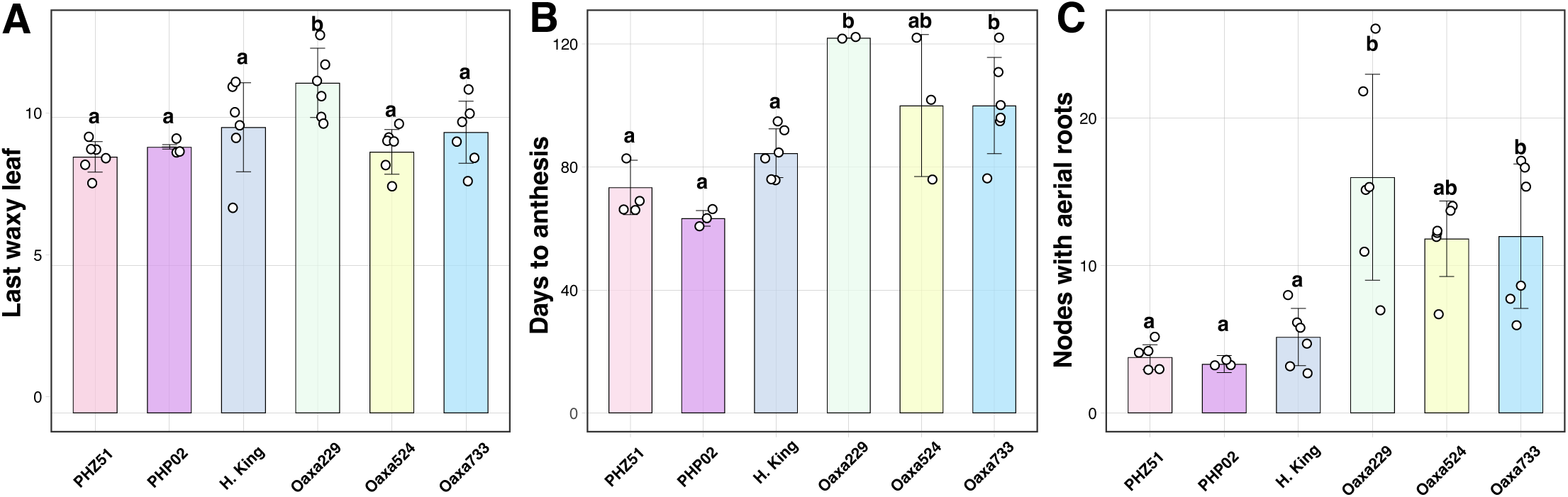
Comparison of exPVP, heirloom, and landrace genotypes in a greenhouse setting. A) Last leaf with epicuticular wax, GH. B) Days to anthesis, GH. C) Number of nodes bearing aerial roots at anthesis. The number of biological replicates was between three and six. ANOVA test was performed with the package multcompView (Ver. 0.1 – 8).

**Figure 3.**
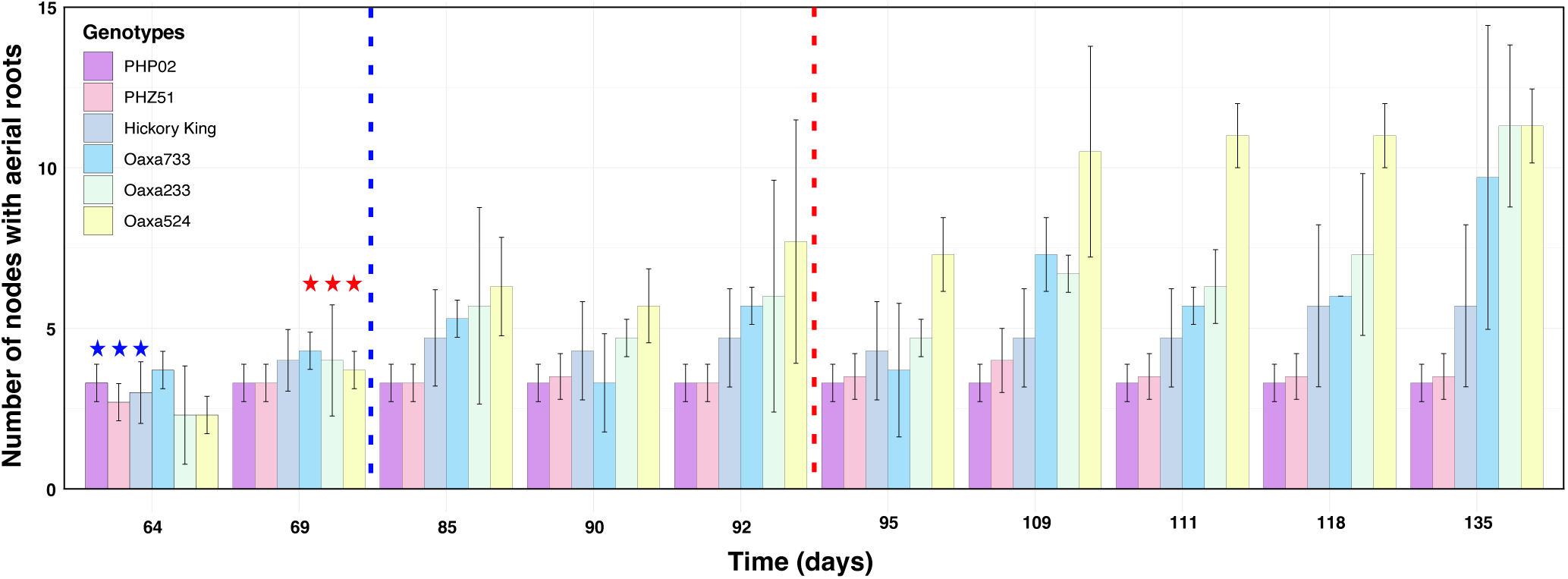
Quantification of aerial roots across six maize genotypes over time. Transitions from juvenile to reproductive and tasseling days are displayed. The bar graph depicts the count of nodes with aerial roots, revealing that landraces exhibit a significantly higher number than exPVP varieties. The Blue dotted line indicated the tasseling of conventional maize lines (70 - 85 DAP), and the red dotted line indicated the tasseling of landraces (95 - 100 DAP). Start indicates the transition from juvenile to adult vegetative stage in landraces.

### Sierra Mixe maize landraces exhibit significant aerial roots during the adult vegetative and reproductive stages in Wisconsin and Georgia under field conditions

#### 2021 field experiment in Wisconsin

To observe landrace and exPVP maize growth-stage-associated traits under field conditions, a trial was established at the West Madison Agricultural Research station in Madison, Wisconsin (WI; summer 2021). Three exPVP varieties (PHZ51, PHP02, and HB8229) and three landrace accessions (Oaxa139, Oaxa524, and GRIN19897) were grown. These landraces were selected from previous studies and field observations in nitrogen fixation in maize (Van Deynze et al., 2018). The exPVPs were selected as material adapted to the Midwest in the US (Mikel, 2006; Van Deynze et al., 2018). Significant differences (p = 0.0395) were found for the last waxy leaf between genotypes (**Supplementary Figure 3A**); however, similar to the greenhouse results (**Figure 2A**), there were no clear patterns differentiating the exPVP genotypes and landrace accessions. For example, the last waxy leaf for PHZ51 did not differ significantly from that of Oaxa139 or Oaxa524, nor did the last waxy leaf for HB8229 differ significantly from PHP02 or GRIN19897. Interestingly, the findings regarding the last waxy appear consistent between PHZ51 and Oaxa524, both in greenhouse and field conditions. Days to anthesis were measured to ascertain the length of the vegetative stage under field conditions. Significant differences ranged from less than 60 days for PHP02 and Oaxa139 to over 100 DAP to anthesis for Oaxa524 and GRIN19897 (**Supplementary Figure 3B**). Three distinct days to anthesis categories were apparent. An early-flowering group (PHP02 and Oaxa 139) reached anthesis in less than 60 DAP. The intermediate-flowering group (PHZ51 and HB8229) reached anthesis in under 70 DAP. Finally, the late-flowering group (Oaxa524 and GRIN19897) had a significantly longer adult vegetative phase, reaching anthesis more than one month later, around 100 DAP. The number of nodes with aerial roots, those regions on the stem generating roots without touching the ground, was counted at anthesis. Significant differences were observed among the genotypes (**Supplementary Figure 3C**). The exPVPs had aerial roots on one to two nodes, whereas landraces had roots on two to six nodes. The landrace accessions Oaxa524 and GRIN19897, which formed aerial roots on significantly more nodes, were the same accessions that took substantially longer to reach the adult reproductive stage (**Supplementary Figure 3B, C and Figure 3**). In the field, we also recorded the number of nodes bearing aerial roots weekly from nine weeks (63 days) to 17 weeks (119 days) post-planting (**Supplementary Figure 4**). This period encompassed the adult reproductive phase of the trial, starting from when PHP02 (the earliest flowering genotype) reached anthesis and continuing until most plants had reached physiological maturity. The exPVP lines (PHZ51, PHP02, and HB8229) all ceased forming aerial roots on nodes once they reached reproductive maturity (anthesis), whereas GRIN19897, Oaxa 524, and Oaxa139 formed an additional node with aerial roots post-anthesis. One particularly close comparison is PHP02 and Oaxa139, which flowered at a similar time (58 and 59 DAP, respectively). PHP02 had two nodes with aerial roots at anthesis and did not develop more nodes with aerial roots, whereas Oaxa139 grew an additional node with aerial roots after anthesis. Landrace genotypes GRIN19897 and Oaxa524 had a long adult vegetative phase and developed five to six nodes with aerial roots by anthesis, and Oaxa524 developed one more node after anthesis (average seven nodes).

#### 2022 field experiment in Georgia

A field trial was established at the University of Georgia Iron Horse Plant Sciences Farm (GA; summer 2022) to observe growth-stage-associated traits in a completely different environment. In this study, six landrace accessions, including two from previous WI studies (Oaxa139, Oaxa524) and four other landrace accessions (Oaxa141, Oaxa306, Oaxa612 and Oaxa622) were grown (**Supplementary Figure 5**). Overall, the trait patterns observed in GA were similar to those seen among the landraces in WI. For the last waxy leaf observations in GA, the range was from leaf 6 to leaf 10 on average (**Supplementary Figure 5A**), whereas the range was from leaf 9 to leaf 11 on average in WI (**Supplementary Figure 3A**). Oaxa139 and Oaxa524 were grown in both locations, and the waxy leaf transition for these genotypes occurred at an earlier growth stage in GA than in WI. In WI, a wide range of days to anthesis was observed among the landrace genotypes. Whereas most accessions took from 75 to >90 days to reach anthesis, on average, one accession, Oaxa 306, reached anthesis significantly earlier, at 52 DAP (**Supplementary Fig 5B**). Interestingly, Oaxa139, grown in both GA and WI, reached anthesis in <60 days in WI, yet it took over 90 days to reach anthesis in GA (**Supplementary Figure 3**). Post-anthesis also significantly differed between the landraces and the exPVPs in different environments **(Figure 2** and **Supplementary Figures 3 and 5)**. The number of nodes with aerial roots at tasselling differed among landrace accessions grown in GA and ranged from 1-14 nodes (**Supplementary Figure 5C**). Oaxa139 and Oaxa524 formed aerial roots on more nodes in GA than in WI.

#### 2022 field experiment in Wisconsin

Finally, exPVPs (PHZ51, PHP02, and HB8229) and landraces (Oaxa139, Oaxa524, Oaxa733, and GRIN19897) were planted in Hancock Agricultural Research Station in central WI in 2022. This third field trial aimed to assess trait expression for a second year in Wisconsin. To ensure comparability, this experiment used an identical set of exPVPs and landraces and an additional one from the previous Wisconsin greenhouse experiment (**Figure 2** and **Supplementary Figure 3**). Since this field was located ∼2 hours away from the main campus, the number of nodes with aerial roots was recorded only at three specific times: 71 (before most accessions flowered), 101, and 116 DAP (after most accessions flowered) (**Supplementary Figure 6**). Consistent with earlier findings, Oaxa524 exhibited the highest average number of nodes with aerial roots (>7). These results affirm that particular maize landraces (e.g., Oaxa524) continue to produce nodes with aerial roots regardless of the environmental context.

### Soil nitrogen rates have a limited effect on maize aerial root development

Insufficient nitrogen availability hinders plant growth and development, including several traits in root architecture (Stitt and Krapp, 1999; Anas et al., 2020). A greenhouse experiment was conducted on three Oaxacan landraces to assess how varying nitrogen levels impact the growth of nodes with aerial roots in maize (GRIN19770, Oaxa139, Oaxa229). Three nitrogen concentrations were applied: 70 ppm (low), 120 ppm (medium), and 170 ppm (high). Data on the number of nodes with aerial roots was collected at 80, 90, and 132 DAP in three biological replicates (**Figure 4)**. No significant differences were observed when comparing the means of each nitrogen treatment within each accession and date, except for accession Oaxa 139, which showed a significant difference between high nitrogen and the other two treatments at 80 DAP. Nitrogen levels did not affect traits such as aerial root diameter, stalk diameter, and plant height (**Supplementary Figure 7**). These results indicate that varying levels of available soil nitrogen do not significantly impact the development of traits related to aerial root development in Oaxacan maize landraces.

**Figure 4.**
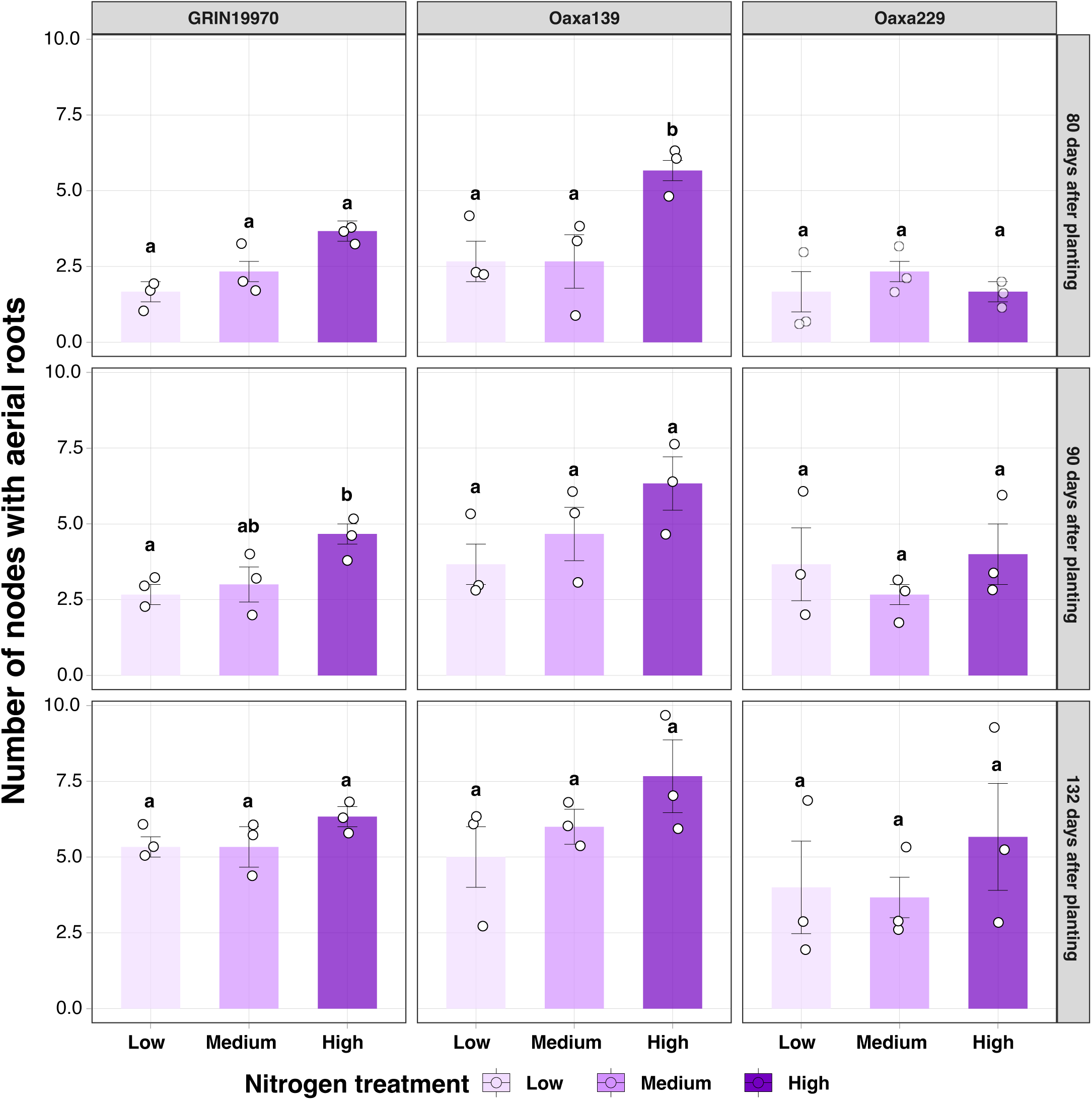
Aerial root development under different nitrogen treatments in maize landraces. The number of nodes with aerial roots was quantified in three landraces (GRIN19770, Oaxa139, and Oaxa229) under three different nitrogen treatments: low (70.36 nitrogen ppm), medium (123.14 nitrogen ppm) and high (175.91 nitrogen ppm). ANOVA was performed using the package multcompView (Ver. 0.1-8) with R (Ver 4.2.1)

### Ambient humidity impacts minimally maize aerial root development

The Oaxacan landraces grow in a high-humidity environment, which enables mucilage secretion from their aerial roots (Van Deynze et al., 2018; Bennett et al., 2020; Pankievicz et al., 2022). Our study aimed to examine how humidity affects the development of aerial roots in the Oaxacan landrace Oaxa524, known for producing numerous nodes with aerial roots under different field and greenhouse conditions. We also used the exPVP PHP02 as a control. We tested two different humidity levels, 30% and 75%. Our experiment focused on measuring the number of nodes with aerial roots, their diameter, and the quantity of aerial roots at the top node. These parameters are associated with mucilage production. (Pankievicz et al., 2022). In the Oaxa524 landrace, there was a slight increase of one node with aerial roots under high humidity (p = 0.021). However, this effect was not observed in PHP02 (**Figure 5A**). Additionally, no significant differences were observed in root diameter or the number of roots at the top node under varying humidity conditions (**Figures 5B and 5C**). These findings indicate a positive but limited effect of humidity in promoting the development of aerial root nodes in this Oaxacan maize landrace.

**Figure 5.**
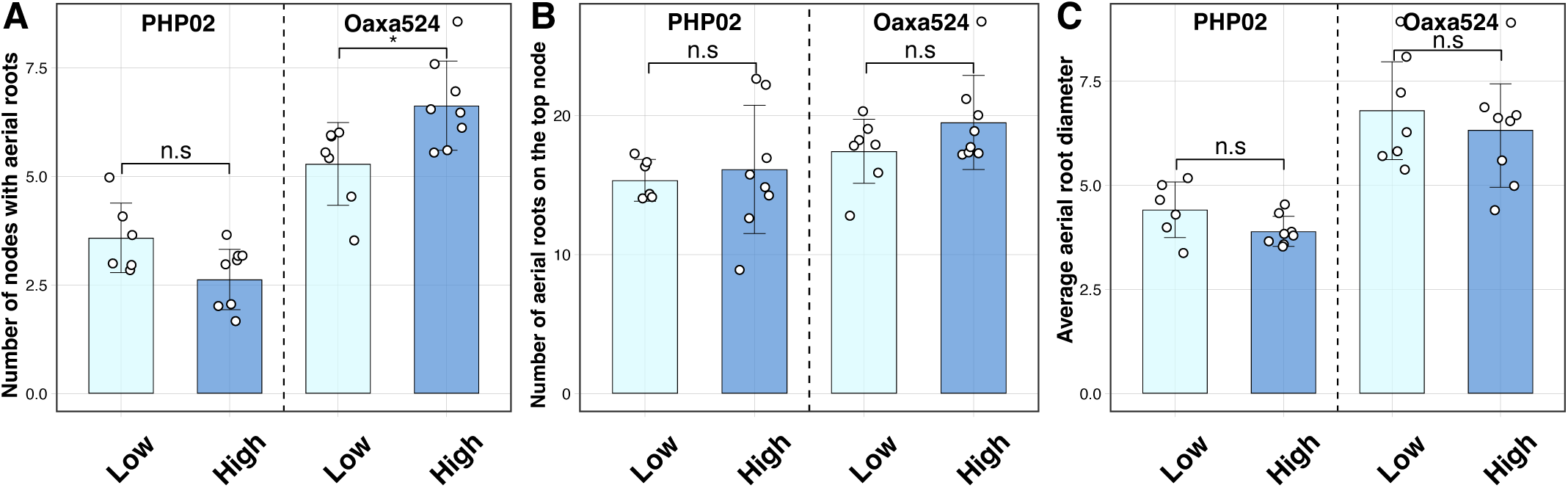
Effect of humidity on aerial root development in an exPVP (PHP02) and an Oaxacan landrace (Oaxa524) accession. Different phenotypes were measured in the maize accessions after flowering. A) number of nodes with aerial roots. B) Number of roots on the top node C) Average aerial root diameter at the top node. A Wilcox test was performed in R (Ver 4.2.1) with the package ggpubr (Ver 0.6.0). Significant levels * p-value < 0.05 and n.s. not substantial.

## DISCUSSION

### Above-ground nodal root development is not a reliable marker for the juvenile stage in tropical maize landraces

The development of above-ground nodal roots (brace and aerial roots) in maize has traditionally been associated with the juvenile stage (Poethig, 1990; Hostetler et al., 2021). Mutants such as *Corngrass1* (*Cg1*) are well known to develop five to six nodes with aerial roots due to an extended juvenile phase as delayed flowering occurs 1 or 2 weeks depending on the inbred background (Poethig, 1988, 2003; Chuck et al., 2007). Numerous maize accessions can develop such roots during the adult stage in response to lodging (Sparks, 2023). However, we showed here that the increased number of nodes with aerial roots in Oaxacan accessions is not due to an extended juvenile phase but to the production of these nodal roots at the adult stage without lodging. Apart from inherent diversity, numerous genes influence nodal root growth, acting as negative regulators (Hostetler et al., 2021). For example, knocking down the *teosinte glume architecture1* (*tga1*) gene led to a rise in the number of nodal roots (Wang et al., 2015). Intriguingly, many of these genes are interconnected with flowering and responses to gibberellic acid (reviewed in (Hostetler et al., 2021)). Nonetheless, flowering in maize is a complex trait governed by many loci with modest impacts (Buckler et al., 2009). This trait is a pivotal adaptation linked to geographical positioning (Ziello et al., 2009; Crimmins et al., 2010). Our observations reveal that the landraces tested here (all cultivated above 1,500 meters) exhibit delayed/late flowering compared to the exPVP lines (**Figure 1A**). Understanding the genetic factors at play in these two traits requires thorough study. Despite research indicating no connection between flowering time and brace root development, it is imperative to investigate the potential presence of genetically correlated variables in these landraces (Reneau et al., 2020). Tropical genotypes are characterized by late flowering, significant height, and increased biomass (leaves) when grown in temperate latitudes (Warrington and Kanemasu, 1983). Hence, delving into these variables is pertinent to gain deeper insights into their suitability for breeding initiatives. We evaluated the epicuticular wax as a morphological marker to determine the transition between juvenile and adult stages. In maize, juvenile leaves have epicuticular wax layers, whereas adult leaves lose this waxy layer and have crenulated epidermal cells (Sylvester et al., 2001). The environment affected this trait, as waxy leaves in Oaxa139 and Oaxa524 genotypes ceased to occur at an earlier growth stage in GA than in WI (**Figure 2** and **Supplementary Figure 4**). In our investigation, through scrutinizing the number of nodes with roots and various traits across three distinct environments, we observed that the landrace Oaxa524 exhibited higher performance compared to other landraces, particularly in terms of number of nodes with aerial roots (**Figures 2, 3,** and **Supplementary Figures 4 and 5**). This insight holds promise for breeding programs incorporating aerial root traits into varieties suited for temperate climates while introducing nitrogen fixation capabilities to reduce reliance on fertilizers. We recognize the necessity for further studies to ascertain the adaptability of these root traits across various environments, spanning from temperate to tropical regions. Our study indicates that above-ground nodal root development should not be used as a marker for the juvenile stage in maize, at least among Oaxacan-like materials.

### Limited effect of soil nitrogen on maize aerial root development

Given the well-known impact of nitrogen limitation on various physiological processes, many farmers globally turn to synthetic fertilizers to boost yields. However, their excessive use poses environmental risks, including water contamination through runoff and economic challenges, especially for small-scale farmers, who often experience the direct effects of price fluctuations influenced by global events (Arndt et al., 2023). With the growing interest in reducing synthetic nitrogen fertilizer use by enhancing nitrogen delivery through biological nitrogen fixation (BNF), it is crucial to understand whether an optimal nutritional rate supports the proper development of aerial roots to maximize the benefits of BNF. Our greenhouse experiment did not reveal any significant effect of soil nitrogen on aerial root development (**Figure 3**). The greenhouse experiment used Turface as the substrate, which offers advantages to control nitrogen levels but differs significantly from field soil, so it would be interesting to perform similar experiments under field conditions in the future.

### Limited effect of ambient humidity on maize aerial root development

Plants display a variety of responses to changes in humidity levels. The biomass of aerial roots in indoor plants is increased due to the high relative humidity within households, facilitated by the indoor environment and misting systems similar to our study (Sheeran and Rasmussen, 2023). Humidity affects the concentration of phytohormones. For instance, cucumbers’ gibberellic acid and auxin levels decrease in response to high humidity (Amin et al., 2021). Similarly, the ability to make more aerial roots was found to be strongly affected by high relative humidity in tomatoes, resulting in a slight reduction in abscisic acid (ABA) levels and an increase in ethylene (Arve and Torre, 2015). Notably, applying exogenous auxin or stem cuttings increased adventitious roots in tomatoes, underscoring the significant influence of hormones and how various stimuli could further enhance this response (Guan et al., 2019). In maize, adventitious roots emerge from aerial nodes, and their development, elongation, and quantity are influenced by a complex interplay of genes and hormones (Singh et al., 2023). Elevated humidity slightly enhanced the number of nodes bearing aerial roots in the landrace Oaxa524 (**Figure 5A**). A phloem-based auxin response may contribute to this increased root formation in maize, akin to what has been observed in rice under saturated humidity (Chhun et al., 2007). Furthermore, ethylene, a plant hormone that accumulates during flooding and induces adventitious nodal roots, could be pivotal in developing nodes with aerial roots in maize. 1-Aminocyclopropane 1-carboxylic acid, which serves as an ethylene precursor, stimulates the generation of adventitious nodal roots in maize (Singh et al., 2023). Consequently, hormone precursors or their inhibitors emerge as a valuable approach to assess the progression of aerial root development in cereal crops. However, humidity did not impact the diameter or the number of roots in our maize accessions (**Figures 5B and C**). In contrast, a study of above-ground roots within the Araceae family showed that the root diameter increases under high humidity conditions (Sheeran and Rasmussen, 2023). Our results suggest that signals such as phytohormones may govern aerial root elongation in Oaxacan maize landraces at the adult stage without any lodging stimulation. Moreover, our study indicated the limited impact of soil nitrogen and ambient humidity on aerial root development, which may facilitate using this trait across diverse environments.

### Implications for biological nitrogen fixation in maize

Our long-term goal is to increase maize’s ability to acquire nitrogen through BNF and thus reduce reliance on exogenous fertilizers. The finding that Oaxacan landraces continue to produce aerial roots into the adult stage has important consequences for breeding since this trait could be introgressed into elite varieties without the potential negative impacts of an extended juvenile phase, such as delayed flowering. The minimal but statistically significant Genotype-by-Enviroment interaction we observed among field sites implies some environmental component to this trait, which may make some environments (e.g., arid ones) unsuitable for trait expression or at least suboptimal. Further research on aerial root production may help identify environment-independent alleles that could be used for breeding. Identifying the specific genes involved would allow for more targeted allele mining or mutagenesis to generate desired traits. Considering the extensive scale of global maize production, even a small reduction in nitrogen requirements could have significant economic and environmental impacts worldwide.

## MATERIALS AND METHODS

### Greenhouse study on aerial root development

In the winter of 2022 at the Walnut Street Greenhouse (Madison, WI), two exPVP (PHZ51, PHP02), one heirloom (Hickory King), and three landraces (Oaxa233, Oaxa524, Oaxa733) were planted in a randomized design with six replications. The greenhouse temperature conditions were established as 25°C during the day, 20°C at night, and a 12-hour light cycle from 6 am to 6 pm was maintained. Plants were irrigated using an automated drip irrigation system three days per week. The last leaf with epicuticular wax was scored on each plant at the V10-V12 stage (**Figure 3A**) following the protocol described by Foerster, 2013; days to anthesis were summed from the planting date to 50% anther dehiscence (**Figure 3B**); the number of nodes with aerial roots longer than 1 centimeter was counted on each plant at anthesis (**Figure 3C**) (Foerster, 2013).

### Wisconsin field studies on aerial root development

In the summer of 2021 at the West Madison Agricultural Research Station (Wisconsin), three exPVP (PHZ51, PHP02, HB8229) and three landrace accessions (Oaxa139, Oaxa524, Oaxa612/GRIN19897) were planted in single-row plots in a randomized complete block design with three replicates. The last leaf with epicuticular wax was scored on three plants per plot at the V10-V12 stage (**Figure 3D**); days to anthesis were summed from planting date to anther dehiscence in 50% of tassels in a plot (**Figure 3E**); the number of nodes with aerial roots was counted on three plants per plot at anthesis (**Figure 3F**). Additionally, the number of nodes with aerial roots was recorded weekly on three plants per plot over nine weeks from 63 DAP to 119 DAP (**Figure 4**). In the summer of 2022 at the Hancock Agricultural Research Station in central WI, three exPVPs (PHZ51, PHP02, and HB8229) and four landraces (Oaxa139, Oaxa524, Oaxa733, and GRIN19897) were planted in single-row plots in a randomized complete block design with three plot replicates and per replicate tree independent plant were measured. The number of nodes with aerial roots was recorded at 71, 101, and 116 DAP.

### Georgia field study on aerial root development

In the summer of 2022 at the University of Georgia’s Iron Horse Farm (Georgia), 11 landrace accessions (Oaxa233, Oaxa229, Oaxa306, Oaxa310, Oaxa141, Oaxa139, Oaxa524, Oaxa612, GRIN19970, Oaxa622, Oaxa182) were planted and screened in single-row plots organized by genotype in a complete block design with three replicates. The phenotypes evaluated during the Wisconsin field study 2021 were also assessed, although the aerial root phenotypes were checked on five plants per plot.

### Greenhouse study on soil nitrogen rates

Three maize accessions (GRIN19770, Oaxa139, and Oaxa229) were planted at the University of Wisconsin-Madison’s Walnut greenhouse (Madison, WI), under controlled conditions of 25°C during the day, 20°C at night, and a 12-hour light cycle from 6 am to 6 pm. Due to the light-sensitive nature of these maize accessions, all windows and walls were covered with black tarps. Seven-gallon pots (Classic 2800) were filled with a mixture of autoclaved calcinated clay, also known as turface (Turface Athletics MVP), and sand in a 4:1 v/v ratio adapted from (Kottkamp et al., 2010). The high temperatures during calcination cause the clay to expand, creating a porous structure. This structure offers physical and chemical stability, providing aeration to plant roots while retaining water within its pore network. This substrate selection aimed to minimize soil potting media variability, ensuring the plants’ consistent growing conditions. Each pot was positioned over a 13-inch plastic pot saucer to prevent the loss of fertilization treatment through the pots’ drainage holes. These trays served as a means to ensure that all the plants had access to the same amount of nitrogen treatment. To investigate the effect of nitrogen fertilization on the number of nodes with aerial roots, three different nitrogen rates were tested: 175 ppm (high), 120 ppm (medium), and 70 ppm (low). These were achieved using various ratios of Hoagland’s solutions with and without nitrogen (HOP01 and HOP03 from Caisson labs). Each treatment comprised three replicates, resulting in a total of 27 pots that were randomly distributed across the greenhouse room. Plants were watered five days a week, receiving 500 ml of the corresponding nitrogen treatment twice a week. The number of nodes with aerial roots was assessed thrice during the experiment at 80, 90, and 132 DAP. Aerial roots were counted as such only if they were not touching the ground and measured 1 cm or more long. In contrast, the root diameter of the aerial roots was measured once at 132 DAP, and three roots were selected randomly from the fourth node (or the highest node if no roots were present in the fourth node).

### Greenhouse study on ambient humidity levels

The maize genotypes Oaxa524 and PHP02 were cultivated in two rooms at the Walnut Street Greenhouses at the University of Wisconsin-Madison. They were grown in 2,800-sized pots (7 gallons) filled with Pro-Mix LP15 medium, with daytime temperatures maintained at 28°C and nighttime temperatures at 25°C. In the greenhouse environment, these plants were exposed to a 12-hour light photoperiod from 6 am to 6 pm and 12 hours of darkness. A humidifier system (Smart Fog, Reno, Nevada) was employed to regulate humidity levels, ensuring a high humidity of 75% and a low humidity of 30%. The plants were irrigated with municipal water twice a week and received fertilization with NPK (20-10-20) weekly. At the flowering stage, the plants were phenotyped for three parameters: the number of nodes with aerial roots, the number of roots on the top node, and aerial root diameter. These measurements were carried out using the established procedure as previously described.

## SUPPLEMENTARY FIGURES

**Supplementary Figure 1.**
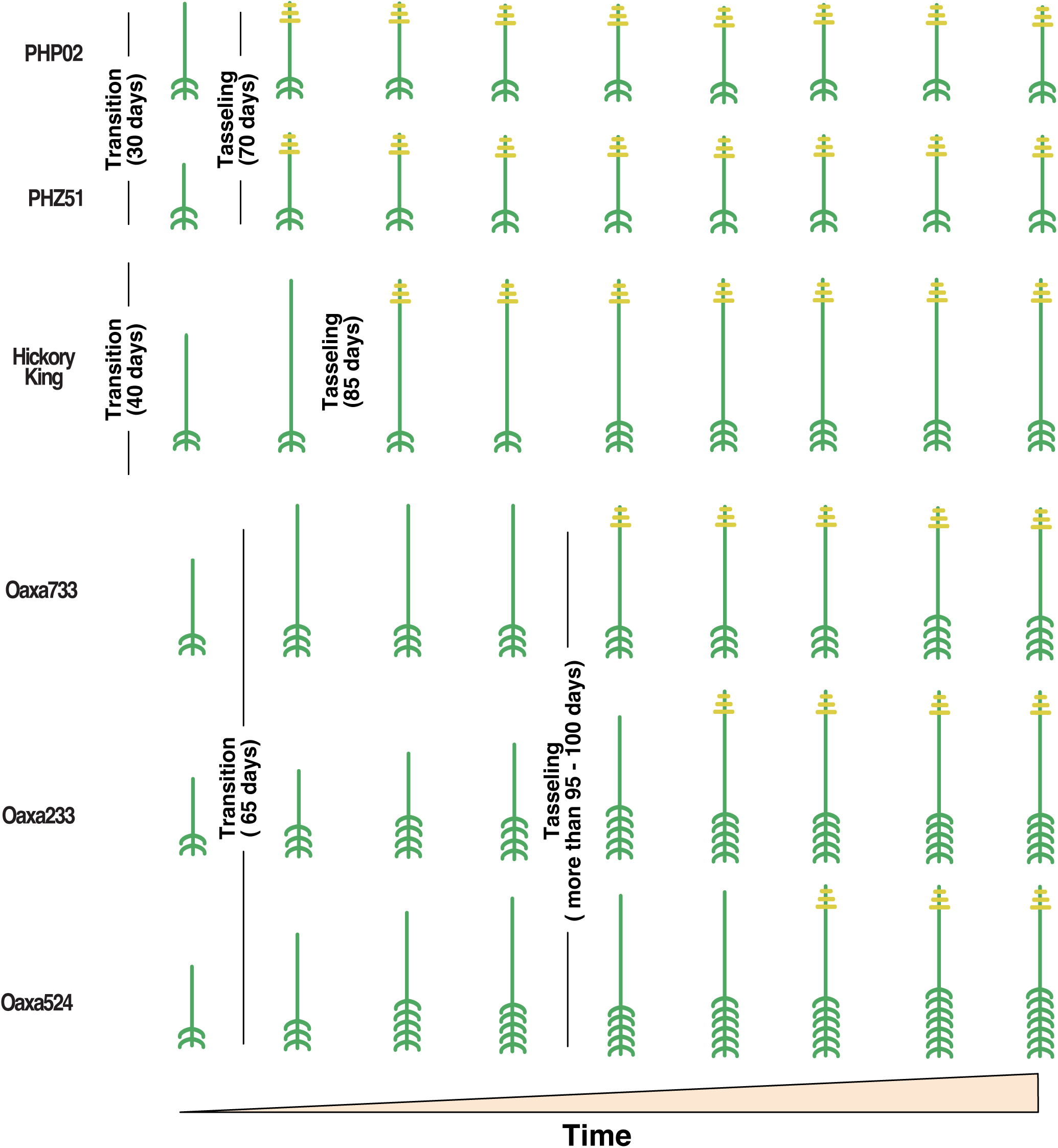
Quantification of aerial roots across six maize genotypes over time. A graphical representation demonstrates the developmental pattern of aerial roots (green) and tasseling (yellow) among different genotypes. Landraces develop aerial roots even after tasseling (reproductive stage), while exPVP varieties cease production upon reaching the adult stage. Transitions from juvenile to reproductive and tasseling days are displayed.

**Supplementary Figure 2.**
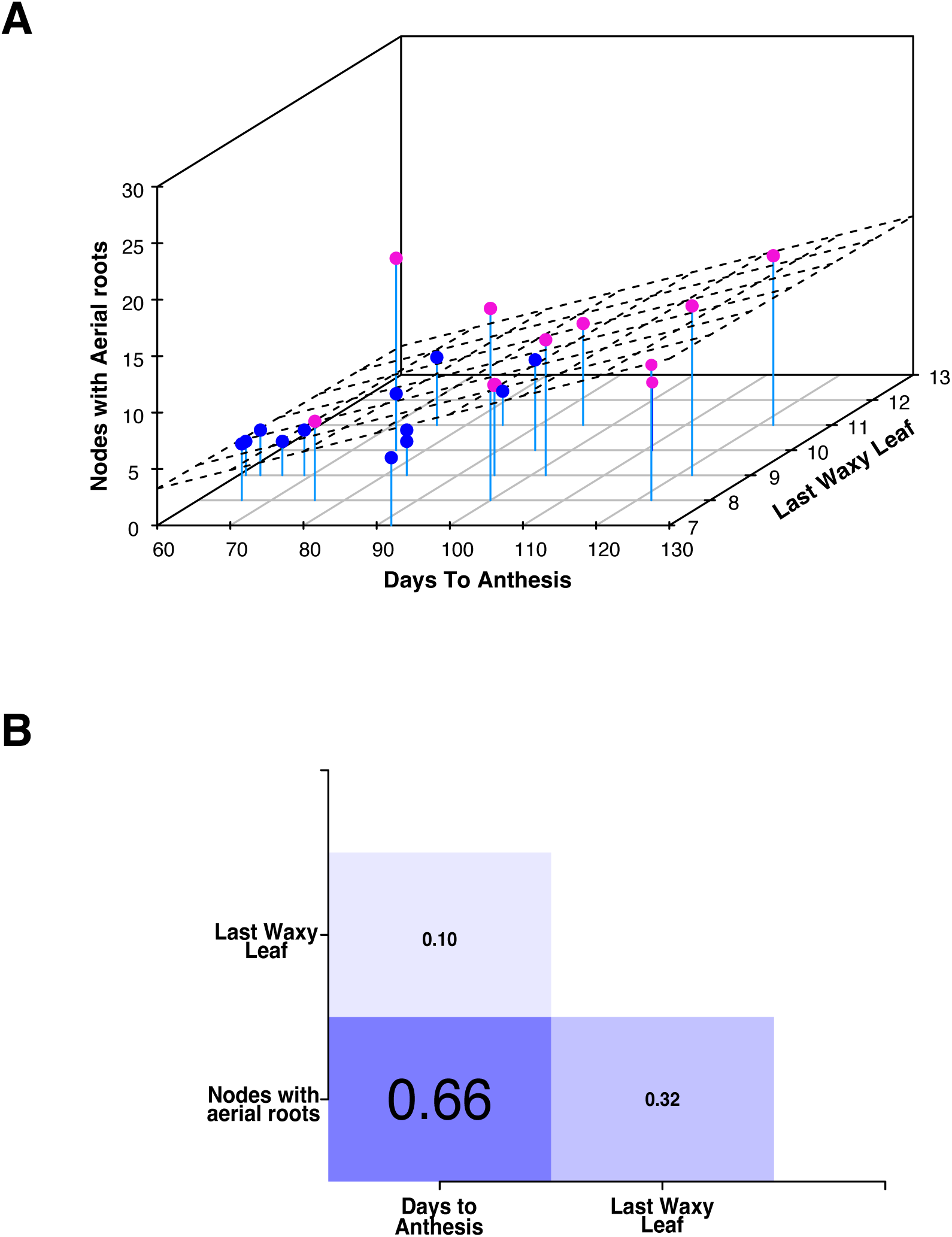
Correlation plots. A) Scatter plot shows the correlation between the number of nodes with aerial roots, days to anthesis, and last waxy leaf. A 3D regression line is included for visualization of the correlation. Landraces (pink dots), exPVP, and Hickory King (blue dots) are depicted. B) Correlation matrix for number of nodes with aerial roots, days to anthesis, Last waxy leaf. Medium to low positive correlations are observed.

**Supplementary Figure 3.**
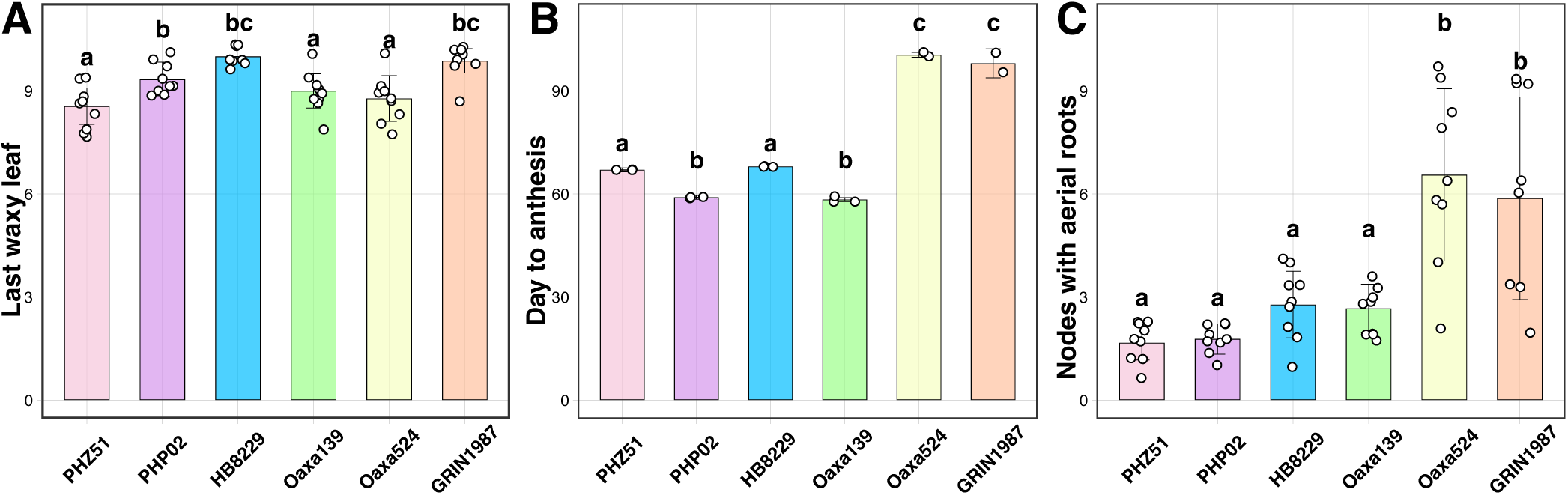
Comparison of exPVP, heirloom, and landrace genotypes in a field study (Wisconsin – WI). A) Last leaf with epicuticular wax. B) Days to tassel anthesis. C) Number of nodes bearing aerial roots at anthesis. ANOVA test was performed with the package multcompView (Ver. 0.1 – 8).

**Supplementary Figure 4.**
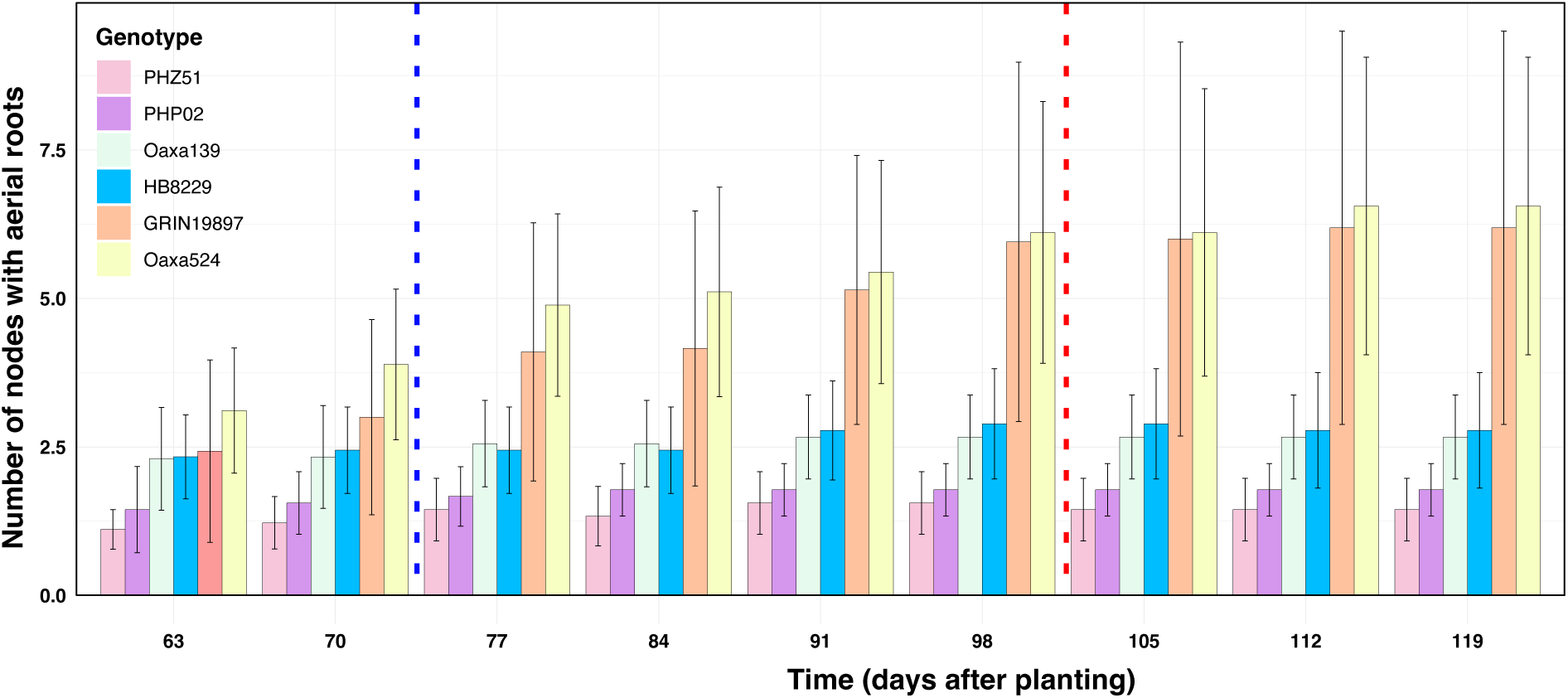
Quantification of aerial roots across six maize genotypes in the field - West Madison Agricultural Research Station 2021. The bar graph depicts the count of nodes with aerial roots. Oaxa524 and GRIN19897 have the most nodes with aerial roots.

**Supplementary Figure 5.**
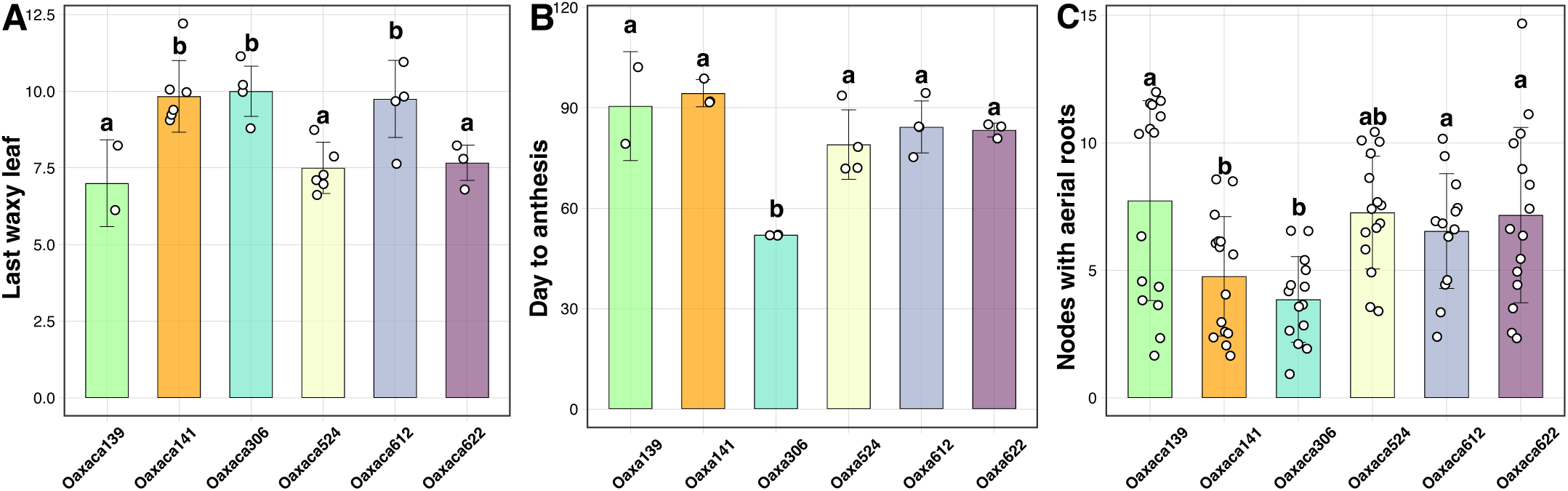
Comparison of different landraces in the field (Georgia – GA). A) Last leaf with epicuticular wax. B) Days to tassel anthesis. C) Number of nodes bearing aerial roots at anthesis. ANOVA test was performed with the package multcompView (Ver. 0.1 – 8).

**Supplementary Figure 6.**
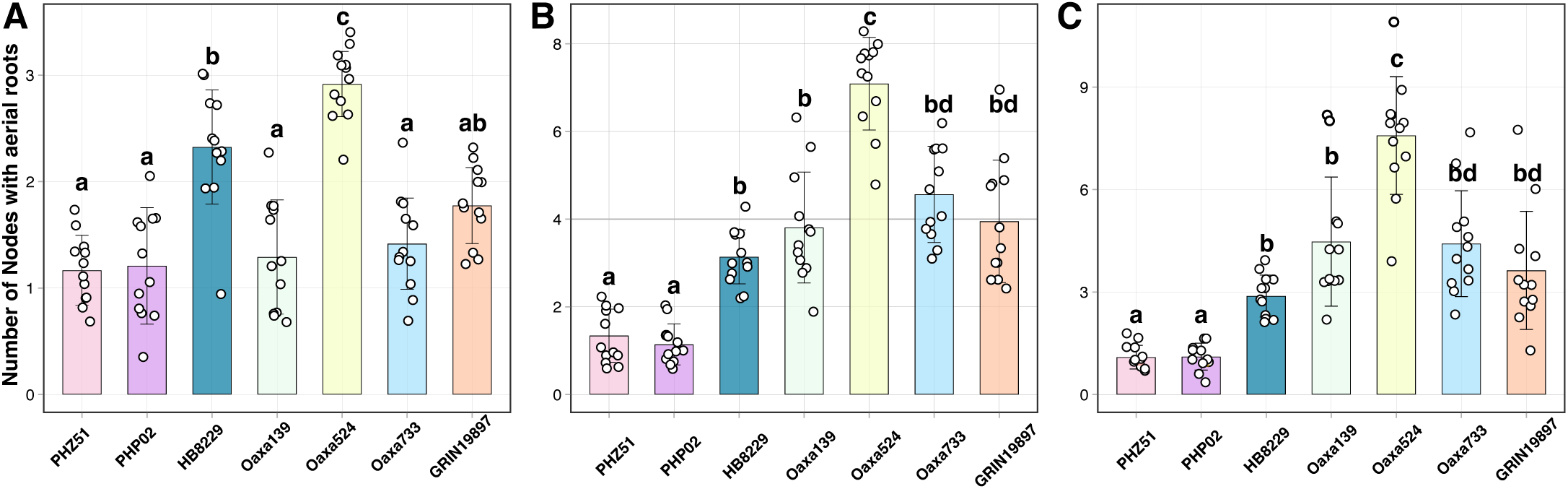
Number of nodes with aerial roots - Field study (Hancock Agricultural Research Station). Each dot is the average of three independent plants per plot. Landraces keep developing nodes with aerial roots compared with exPVP lines. A) Early measurement 71 DAP, B) Mid-season measurement 101 DAP. C) Late-season measurement 116 DAP. ANOVA test was performed with the package multcompView (Ver.0.1 – 8) and n = 12

**Supplementary Figure 7.**
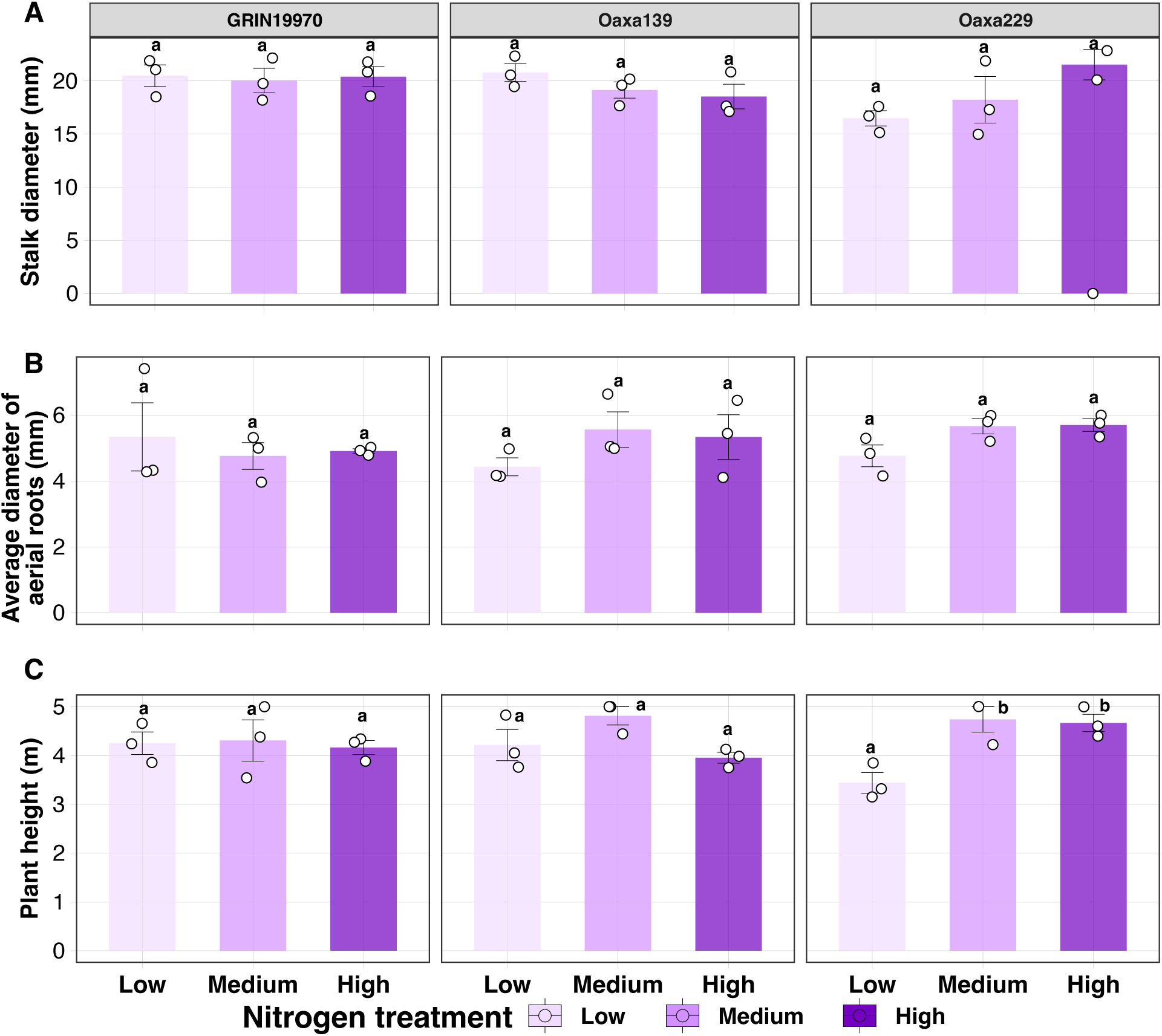
Additional traits were measured in the greenhouse under different nitrogen treatments in maize landraces. Three additional traits were measured in the landraces: GRIN19770, Oaxa139, and Oaxa229 under three different nitrogen treatments: low (70.36 nitrogen ppm), medium (123.14 nitrogen ppm), and high (175.91 nitrogen ppm). These traits included stalk diameter, the average diameter of aerial roots, and plant height. ANOVA was performed using the package multcompView (Ver. 0.1-8) with R (Ver 4.2.1)

