## Supplementary Figures for "Aerial root formation in Oaxacan maize (*Zea mays*) landraces persists into the adult phase and is minimally affected by soil nitrogen and ambient humidity"

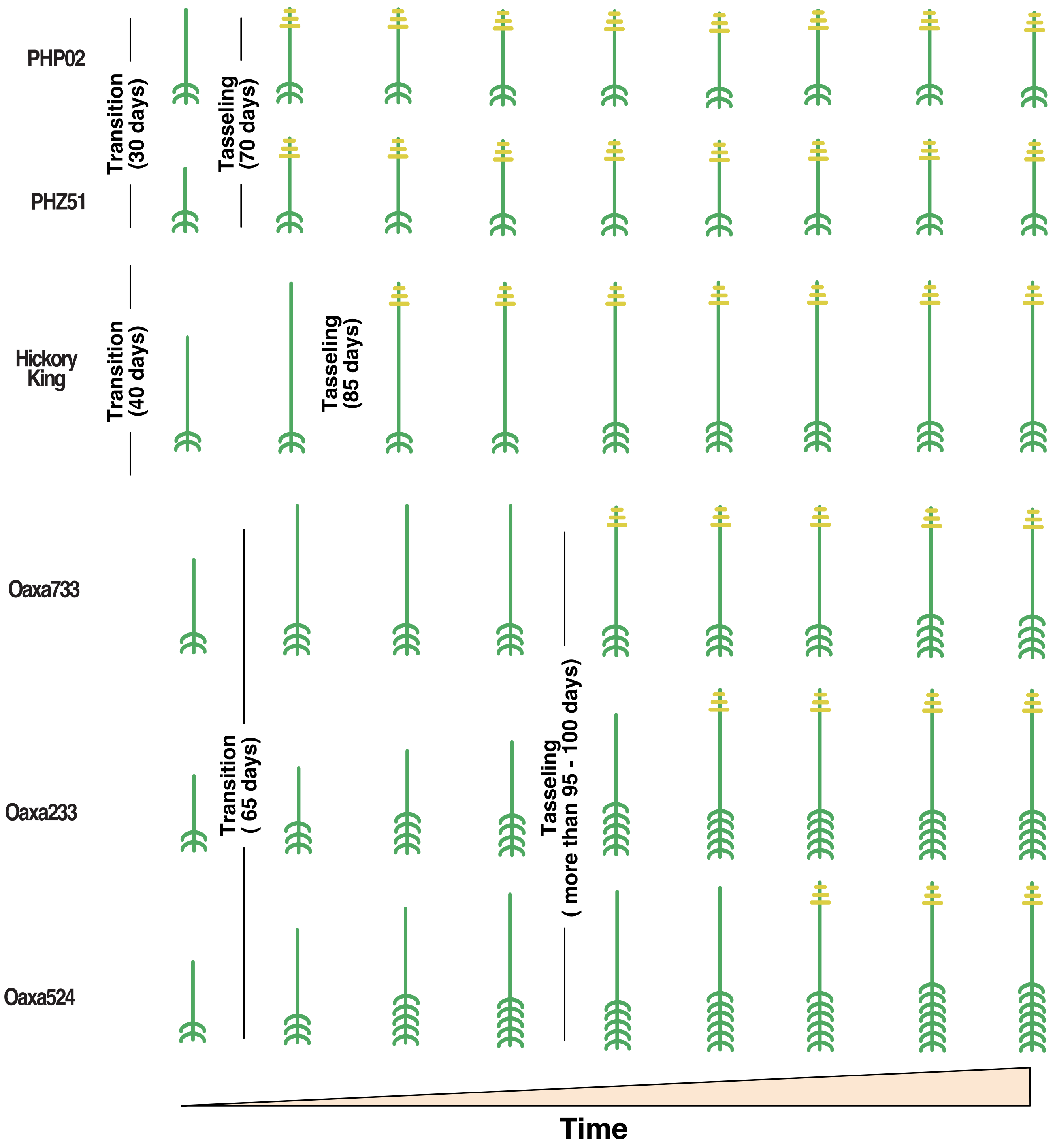


**Supplementary Figure 1.** Quantification of aerial roots across six maize genotypes over time. A graphical representation demonstrates the developmental pattern of aerial roots (green) and tasseling (yellow) among different genotypes. Landraces develop aerial roots even after tasseling (reproductive stage), while exPVP varieties cease production. Transitions from juvenile to reproductive and tasseling days are displayed.

A

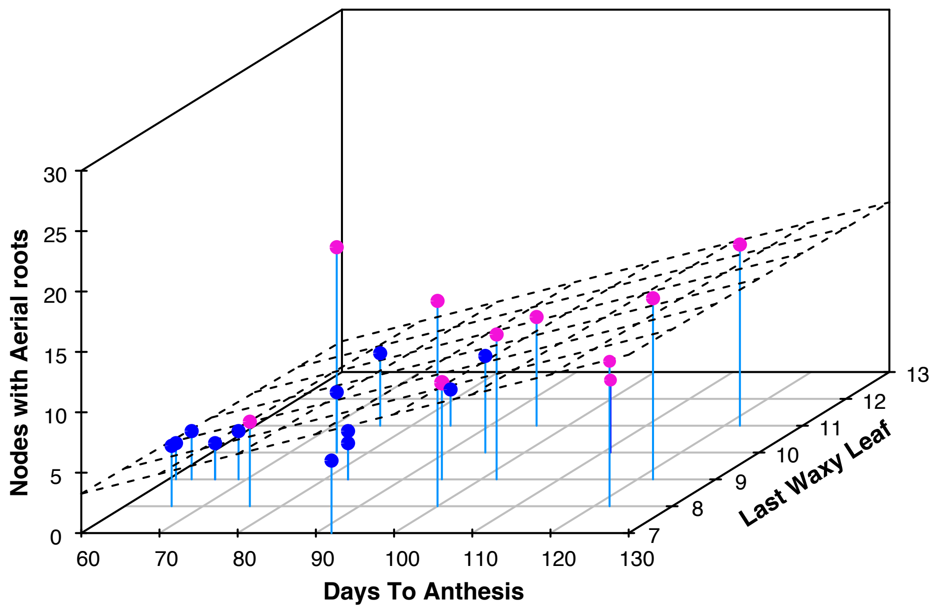


B

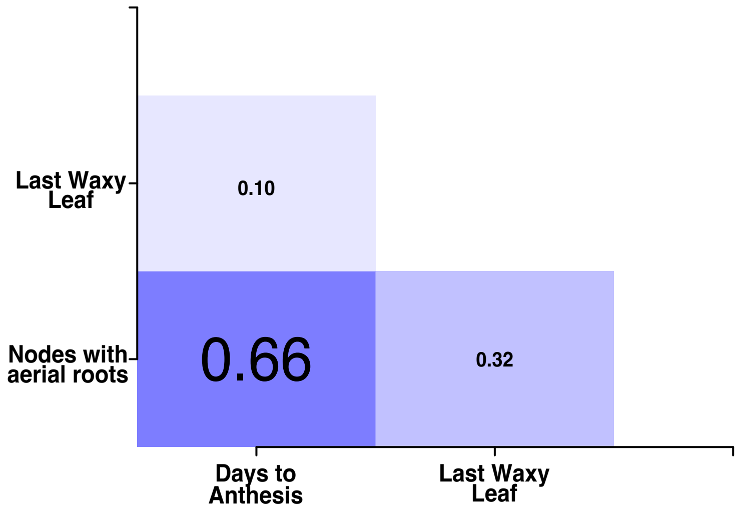


**Supplementary Figure 2. Correlation plots.** A) Scatter plot shows the correlation between the number of nodes with aerial roots, day to anthesis, and last waxy leaf. A 3D regression line is including for visualization of the correlation. Both landraces (pink dots) and exPVP (blue dots) are depicted. B) Correlation matrix for number of nodes with aerial roots, days to anthesis, Last waxy leaf. Medium to low positive correlations are observed.

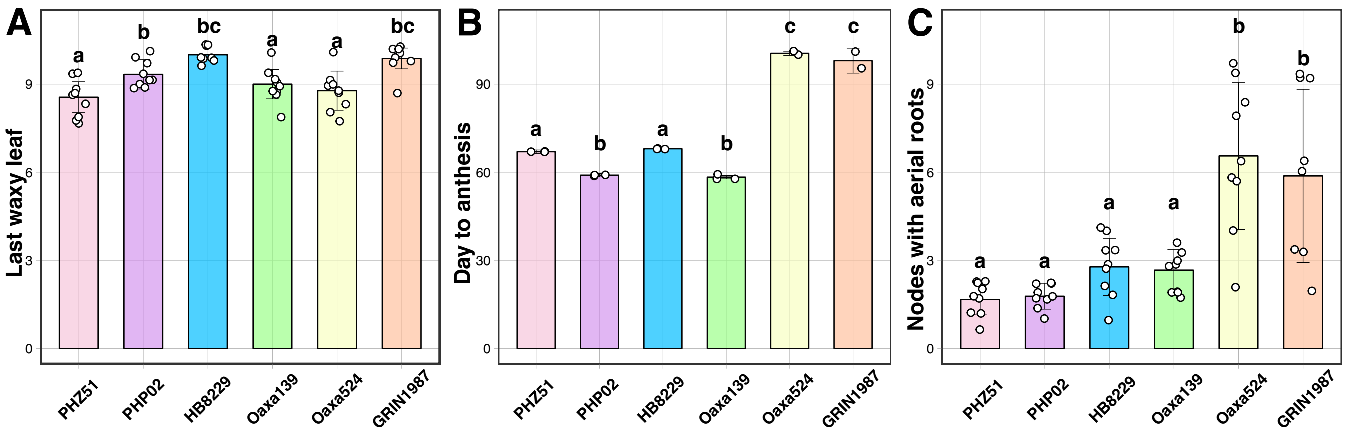


**Supplementary Figure 3.** Comparison of exPVP, heirloom and landrace genotypes in a field study (Wisconsin – WI). A) Last leaf with epicuticular wax. B) Days to tassel anthesis. C) Number of nodes bearing aerial roots at anthesis. Number of biological replicates between three to nine. ANOVA test was performed with the package multcompView (Ver. 0.1 – 8).

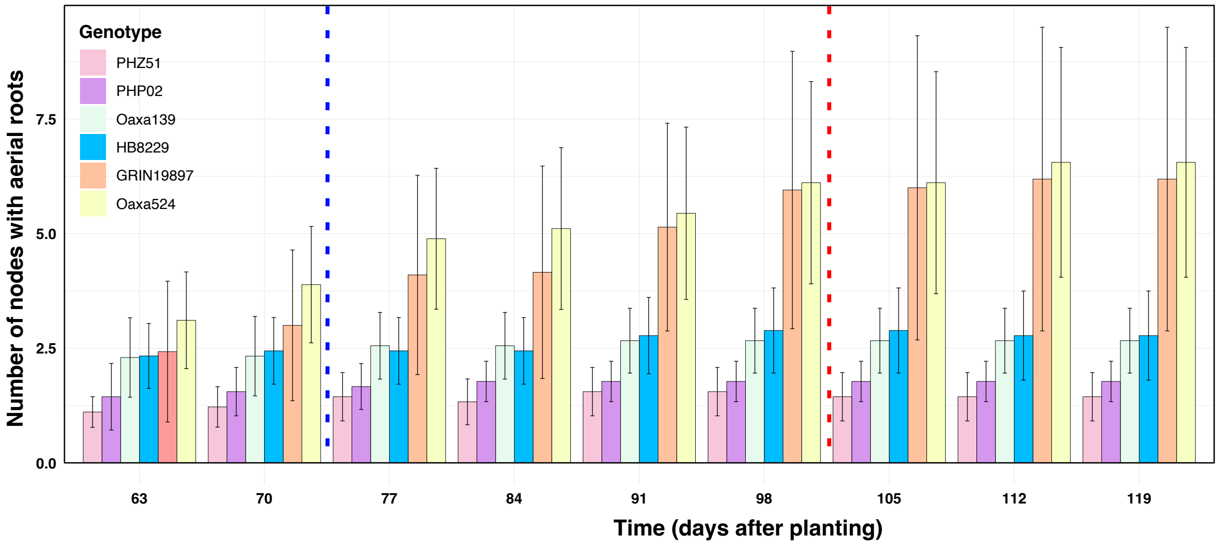


**Supplementary Figure 4.** Quantification of aerial roots across six maize genotypes in the field - West Madison Agricultural Research Station 2021. The bar graph depicts the count of nodes with aerial roots. Oaxa524 and GRIN19897 have the most nodes with aerial roots.

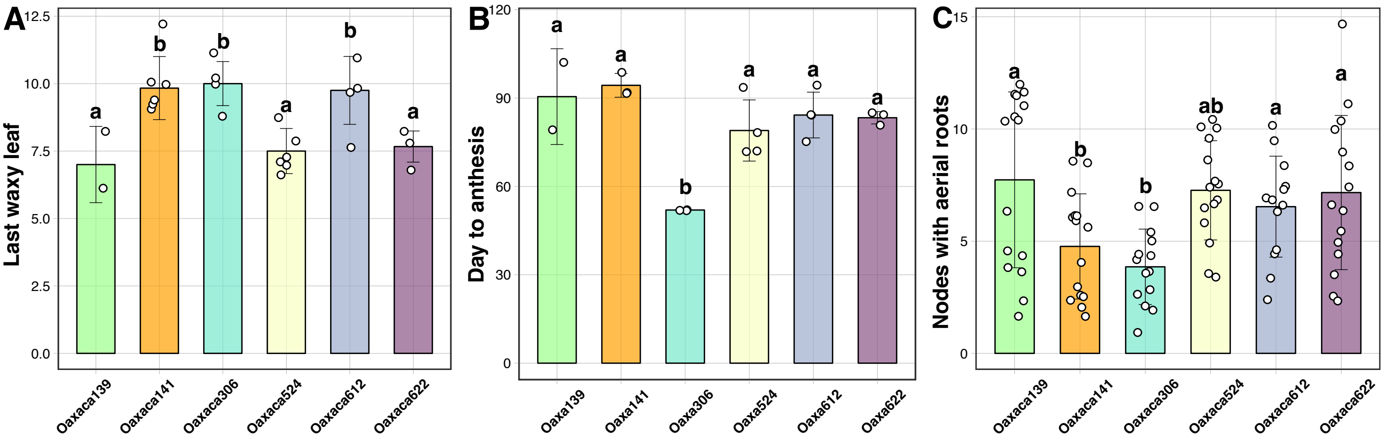


**Supplementary Figure 5.** Comparison of different landrace in the field (Georgia – GA). A) Last leaf with epicuticular wax. B) Days to tassel anthesis. C) Number of nodes bearing aerial roots at anthesis. ANOVA test was performed with the package multcompView (Ver. 0.1 – 8).

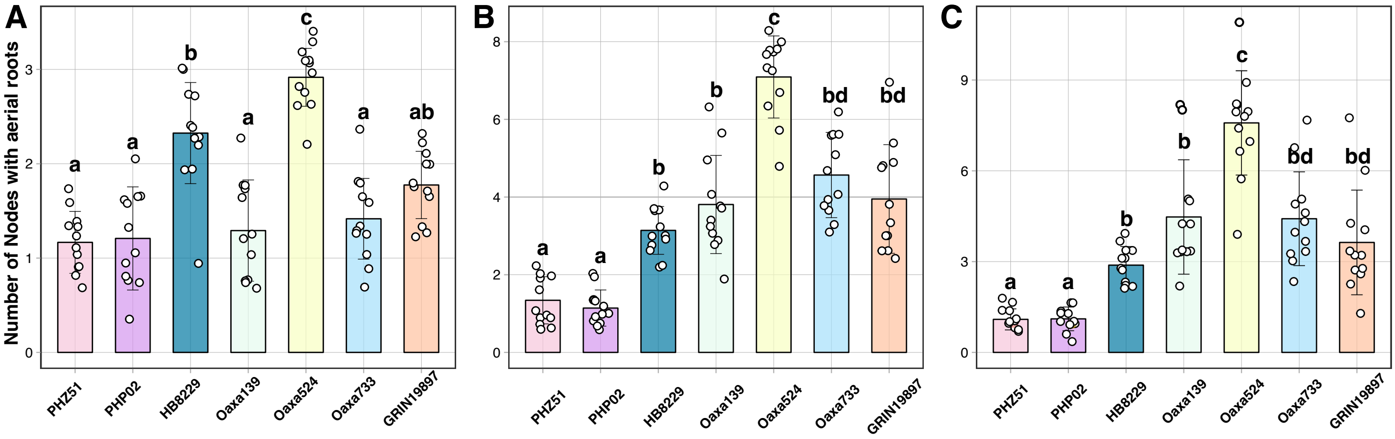


**Supplementary Figure 6.** Number of nodes with aerial roots - Field study (Hancock Agricultural Research Station). Each dot is the average of three independent plants per plot. Landraces keep developing nodes with aerial roots compared with exPVP lines. A) Early-season measurement 71 DAP, B) Mid-season measurement 101 DAP. C) Early measurement 116 DAP. ANOVA test was performed with the package multcompView (Ver. 0.1 – 8) and n = 12

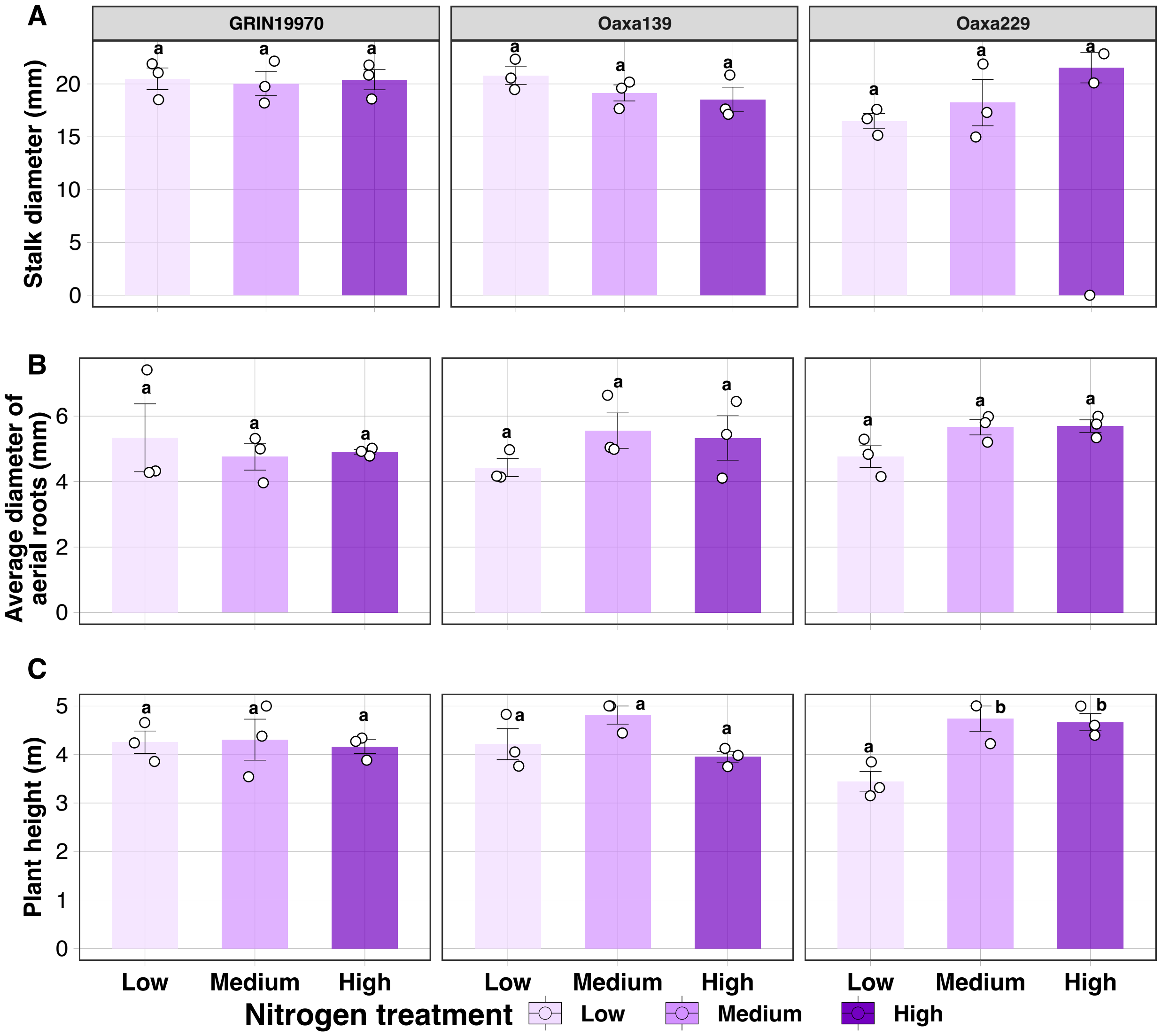


**Supplementary Figure 7. Additional traits under different nitrogen treatments in maize landraces.** Three additional traits were measured in the landraces: GRIN19770, Oaxa139 and Oaxa229 under three different nitrogen treatments low (70.36 nitrogen ppm), medium (123.14 nitrogen ppm) and high (175.91 nitrogen ppm). These traits included stalk diameter, average diameter of aerial roots and plant height. ANOVA was performed using the package multcompView (Ver. 0.1-8) with R (Ver 4.2.1)
